# Cellograph: A Semi-supervised Approach to Analyzing Multi-condition Single-cell RNA Sequencing Data Using Graph Neural Networks

**DOI:** 10.1101/2023.02.24.528672

**Authors:** Jamshaid A. Shahir, Natalie Stanley, Jeremy E. Purvis

## Abstract

With the growing number of single-cell datasets collected under more complex experimental conditions, there is an opportunity to leverage single-cell variability to reveal deeper insights into how cells respond to perturbations. Many existing approaches rely on discretizing the data into clusters for differential gene expression (DGE), effectively ironing out any information unveiled by the single-cell variability across cell-types. In addition, DGE often assumes a statistical distribution that, if erroneous, can lead to false positive differentially expressed genes. Here, we present Cellograph: a semi-supervised framework that uses graph neural networks to quantify the effects of perturbations at single-cell granularity. Cellograph not only measures how prototypical cells are of each condition but also learns a latent space that is amenable to interpretable data visualization and clustering. The learned gene weight matrix from training reveals pertinent genes driving the differences between conditions. We demonstrate the utility of our approach on publicly-available datasets including cancer drug therapy, stem cell reprogramming, and organoid differentiation. Cellograph outperforms existing methods for quantifying the effects of experimental perturbations and offers a novel framework to analyze single-cell data using deep learning.

## 1 Introduction

The rapid progression of single-cell technologies (Klein et al., 2015) has enabled scientists to accumulate complex datasets to study differentiation and developmental trajectories in response to differing experimental perturbations, assess the efficacy of a drug in treating of disease, and evaluate the efficiency of different reprogramming protocols. Regardless of the preceding experimental design, many single cell RNA sequencing analyses follow the same pipeline (Luecken and Theis, 2019): pre-processing and quality control followed by clustering and differential gene expression. In the context of studying more continuous phenomena such as differentiation or cell reprogramming, trajectory analysis may also be employed (Haghverdi et al., 2016). However, in the case of multiple experimental conditions such as different time points sampled for sequencing in cell reprogramming, or varying concentrations of a cancer drug, these methods may fall short in faithfully summarizing the underlying biology. In particular, clustering and differential gene expression give a bulk summary of the transcriptomic variation between computationally-inferred discrete populations, but do not explicitly consider the single-cell variability within treatment groups, such as how prototypical an individual cell is of its assigned treatment group.

### 1.1 Related Methods

Differential abundance methods can rectify these challenges by quantifying differences between and within conditions at a finer resolution. Milo tests for differential abundance on *k*-nearest neighbor (kNN) graphs by aggregating cells into overlapping neighborhoods and performing a quasi-likelihood F test. Dann et al. (2021). This returns a metric of the log-fold change of the differential abundance in each neighborhood. However, because Milo aggregates cells into neighborhoods, it does not provide single-cell resolution providing insight into the impact each perturbation has on an individual cell.

Covarying Neighbor Analysis (CNA) Reshef et al. (2021) performs association analysis agnostic of parameter tuning, making it an efficient method. Like Milo, it aggregates cells into neighborhoods, and calculates a neighborhood abundance matrix (NAM), where each entry *C_n_, m* is the relative abundance of cells from sample *n* in neighborhood *m*. From there, it derives principal components where positive loadings correspond to higher abundance while negative loadings correspond to lower abundance. This enables the characterization of transcriptional changes corresponding to maximal variation in neighborhood abundance across samples. Association testing is performed between transcriptional changes and attributes of interest using the first *k* NAM-PCs. It returns the Spearman correlation between the attribute and abundance of the neighborhood anchored at each cell, providing a single-cell metric. However, its performance falls short when considering more than two conditions.

MELD (Burkhardt et al., 2021) sought out to quantify the effect of an experimental perturbation on individual cells in scRNA-seq data using graph signal-processing to infer a sample-associated density that is then normalized to give a probability of each cell belonging to a condition of interest defined as a relative likelihood. The authors also introduced a novel clustering approach, called Vertex Frequency Clustering (VFC), which clusters data according to not just transcriptomic similarity but also how the MELD-derived relative likelihood scores, thereby identifying populations of cells similarly enriched or depleted in conditions according to the perturbation response. However, the original study restricted evaluation to datasets with two conditions to discriminate between: a control condition and a single perturbed condition, and therefore did not consider multiple treatment conditions, which are prevalent and can provide more insight, for instance, the response of a drug at various time intervals, combining drugs, or administration of a differentiation stimulus at different time-points. Furthermore, robust calculation of the sample-associated likelihood relies on computationally-expensive parameter estimation that can take upwards of 12 hours with 36 cores on a high-performance computing cluster for a dataset of 26,827 cells. In addition, VFC is memory-intensive, which limits its scalability to larger datasets.

### 1.2 Graph Neural Networks

In recent years, the rapidly-emerging field of deep learning has seen utility in scRNA-seq analysis (Amodio et al.,2019; Lopez et al., 2018; Wang et al., 2021). More recently, graph neural networks (GNN) have demonstrated promise in capturing the structural information of scRNA-seq data via the graphical representation the high-dimensional assay naturally lends itself towards, with cells as vertices or nodes, and edges between them representing similarity in gene expression. This connectivity enables the model to naturally leverage the relationship between similar cells in a variety of tasks, most notably clustering and imputation. GNNs pass in graphical representations of data as input to perform a myriad of classification tasks, namely node classification, edge classification, and graph classification. Unlike convolutional neural networks (CNNS) which involve multiple layers and can take a long time to train depending on the size of the data, GNNs require only a few layers to achieve high performance in a fraction of the time. Furthermore, whereas CNNs require large amounts of training data, GNNs can learn patterns in data in a semi-supervised fashion: they take the entire data structure as input, but only a paucity of nodes are labeled; a larger portion is held out for validation and testing purposes. The applications of GNNs has been demonstrated in the case of graph classification, edge classification, and node classification. For example, scGNN (Wang et al., 2021) used GNNs and a Gaussian mixture model to perform imputation and cell clustering. Another study used Graph Attention Networks (GATs) (Ravindra et al., 2020), (Veličković et al., 2017) which are a subset of GNNs based on the self-attention mechanism commonly used in natural language processing, to predict disease state in scRNA-seq data from multiple sclerosis patients, followed by another study from the same group applied to COVID-19 patients (Sehanobish et al., 2020). GATs have also been used as part of variational graph autoencoders to facilitate clustering. Finally, GNNs have been used in conjunction with relational networks to predict breast cancer subtypes in bulk RNA-seq data (Rhee et al., 2018). However, their potential to ascertain the responsiveness of individual cells to perturbations in order to gauge the efficacy of the experimental stimulus, particularly in complex experimental designs that span multiple conditions or time points, has not been formally assessed.

In this work, we introduce Cellograph: a novel computational framework using Graph Convolutional Networks (GCN) (Kipf and Welling, 2016) - which are the graph analogue of CNNs - to perform node classification on scRNA-seq data collected from multiple conditions, treating the individual cells as nodes. Cellograph uses a two-layer GCN to learn a latent representation of the single-cell data according to how representative each cell is of its ground truth sample label. This latent space can be easily clustered to derive groups of cells associated with similar treatment response and transcriptomics, as well as projected into two dimensions for visualization purposes. Cellograph outperforms existing approaches in quantifying the effects of perturbations and offers a novel GNN framework to cluster and visualize single-cell data. In the following sections, we discuss the workflow of Cellograph, demonstrate its performance on three published scRNA-seq datasets, and benchmark it against previously published methods using cross-categorical entropy and normalized mutual information (McDaid et al., 2011).

## 2 Methods

### 2.1 Overview of the Cellograph algorithm

Consider a data set of *n* cells with *m* genes measured. Cellograph learns a low-dimensional representation of each cell through node classification. A kNN graph representation of the data is first constructed where each cell is represented by the top 10-30 principal components (PCs) and edges connect cells that are similar in this space (Figure 1A). We define this *k*NN graph as a tuple *G* = (*V, E*), where *V* is the set of vertices or nodes, and *E* is the set of edges between nodes. A common matrix representation of *G* is the adjacency matrix 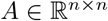. We train a two-layer GCN on the derived graph as follows:

**Figure 1:**
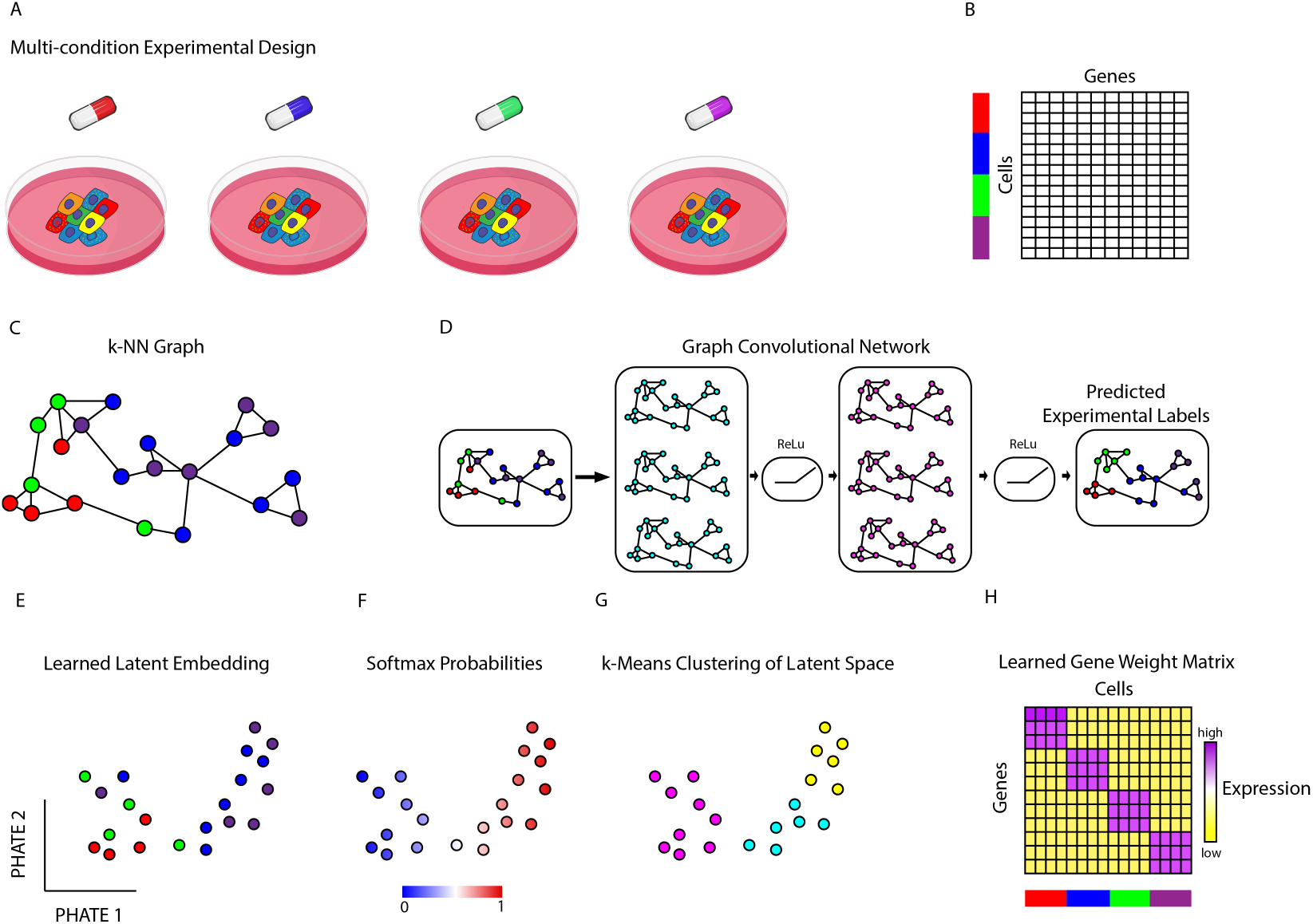
Illustrative overview of Cellograph algorithm. Single-cell data collected from multiple sample drug treatments (A,B) is converted to a kNN graph (C), where cells are nodes, and edges denote connections between transcriptionally similar cells. The colored rectangles (B) correspond to the different samples represented by the drugs in (A). This kNN is fed in as input to a two-layer GCN (D) that quantitatively and visually learns how prototypical each cell is of its experimental label through the learned latent embedding (E) and softmax probabilities (F), respectively. The learned latent space is amenable to k-means clustering (G). Finally, we can use a weight matrix (H) parameterized by the genes in our dataset to identify the most informative genes in distinguishing between conditions.

Considering *c* experimental conditions, let *Z* be an *n* by *c* matrix of probabilities of observing a cell in an experimental condition. We define *Z* as follows:

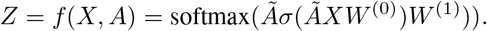

Here, 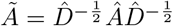 and 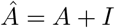.

We define each hidden layer as the normalized graph Laplacian

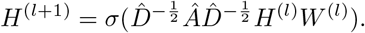

The softmax function is defined as

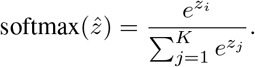

Each 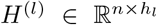 is a latent embedding of the nodes in layer *l*, *Â* = *A* + *I* is the symmetric adjacency matrix, 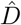 is the degree matrix of *Â*, with degrees of each node along the diagonal, *W*^(l)^ is a learned weight matrix for layer *l*, and *σ* is a LeakyRelu function for GCNs. The GCN learns an embedding 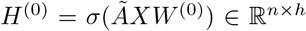 it iteratively orders cells according to how prototypical they are of the experimental labels in question in *h*-dimensional space (in all of our experiments, *h* = 16).

We project cells based on their computed latent space representation into two dimensions with PHATE (Moon et al., 2019), which gives a more interpretable visualization than applying PHATE to the original data by learning a lower-dimensional representation of the data based on both transcriptomic similarity, and similarity across ground truth conditions. We can also directly apply *k*-means clustering to *H*^(0)^, obtaining more informative clusters as evidenced by higher normalized mutual information (NMI) scores (McDaid et al., 2011), a common metric for benchmarking clustering algorithms that measures how concordant the clusters were with the original ground truth sample assignments. *W*^(1)^ is an *h* × *m* learned weight matrix, where m is the number of genes. If we take the transpose of *W*^(1)^ and rank the summed entries across rows, we can identify the top-weighted genes the model found most informative in learning the difference between the experimental conditions. Unlike differential gene expression, which is generally applied to arbitrarily clustered data, the learned weight matrix can provide a direct transcriptomic comparison between ground truth conditions agnostic of statistical models.

As mentioned previously, GNNs learn in a semi-supervised manner, where the entire graph is seen during the training, but only a fraction of the nodes are labeled. We randomly select 1 – 3% of nodes from each condition as training nodes where labels are not hidden, and sample a larger proportion for testing and validation. We employ a categorical cross-entropy loss function for minimization during training. Unless specified otherwise, we train the GNN for 200 epochs with a patience of 30, which is the number of epochs with no improvement after which the training terminates. The source code for Cellograph is available at https://github.com/jashahir/cellograph

## 3 Results

We demonstrated the biological application of Cellograph on three published single-cell RNA sequencing datasets: a human organoid model of intestinal stem cells differentiating to Paneth cells with or without a stimulus to enhance the efficiency of the differentiation (Mead et al., 2022); a non-small-cell lung carcinoma (NSCLC) cell line that was treated with a drug called Erlotinib at various time points and later temporarily withdrawn from the drug for several days (Aissa et al., 2021); and a myogenesis model of transdifferentiation and traditional cell reprogramming (Yagi et al., 2021). The metadata and digital gene expression data for the human organoid dataset was downloaded from https://singlecell.broadinstitute.org (study SCP1318). The Erlotinib drug holiday dataset was downloaded from the National Center for Biotechnology Information’s database Gene Expression Omnibus (GEO) (https://www.ncbi.nlm.nih.gov/geo) under the accession number GSE134841. The myogenesis dataset was downloaded from GEO under the accession number GSE171039. We benchmarked the performance of Cellograph against the aforementioned differential abundance methods, MELD, Milo, and CNA. Our results show robust performance of Cellograph on these distinct datasets, and provide valuable biological insights.

### 3.1 Cellograph captures shifts in cell type abundance during human intestinal organoid differentiation

We first applied Cellograph to an organoid model of intestinal stem cells differentiating to Paneth cells with or without KPT-330, an inhibitor of the nuclear exporter, Exportin 1, which was demonstrated in the original study (Mead et al.,2022) to enhance the abundance of Paneth cells following differentiation. Samples were collected from 6 donors for sequencing following 6 days of treatment with or without KPT-330. We will refer to these two conditions as KPT and control cells, respectively. We trained Cellograph using a two-layer GCN with 80 out of the 2484 cells labeled, such that 40 were labeled for each condition. We projected the learned latent space to 2 dimensions with PHATE and colored cells according to the probability of belonging to the KPT-treated condition (Figure 2A). We obtain a smooth gradient of cells along the PHATE plot, with cells arranged according to how impacted they are by KPT treatment. To determine if this gradient reflected meaningful biology, we extracted the 25 top-weighted genes from the learned weight matrix and visualized them with a heatmap categorized by the two treatment groups (Figure 2B). This is derived from the first layer of the GCN and parameterizes each gene, where the model upweights genes it finds most relevant in distinguishing between conditions. Among the top 25 genes is GDF15, a marker of DUOX2+ WAE-like and WAE-like cells, which is highly expressed in KPT-treated cells, where these cell types are more abundant due to the greater efficiency of Paneth cell differentiation (Zhang et al., 2019; Yang et al., 2021). Conversely, KLK6 is highly expressed in the control-treated population, which has been shown to mediate the multipotency of intestinal stem cells (Schrader et al., 2015; Zhou et al., 2021).

**Figure 2:**
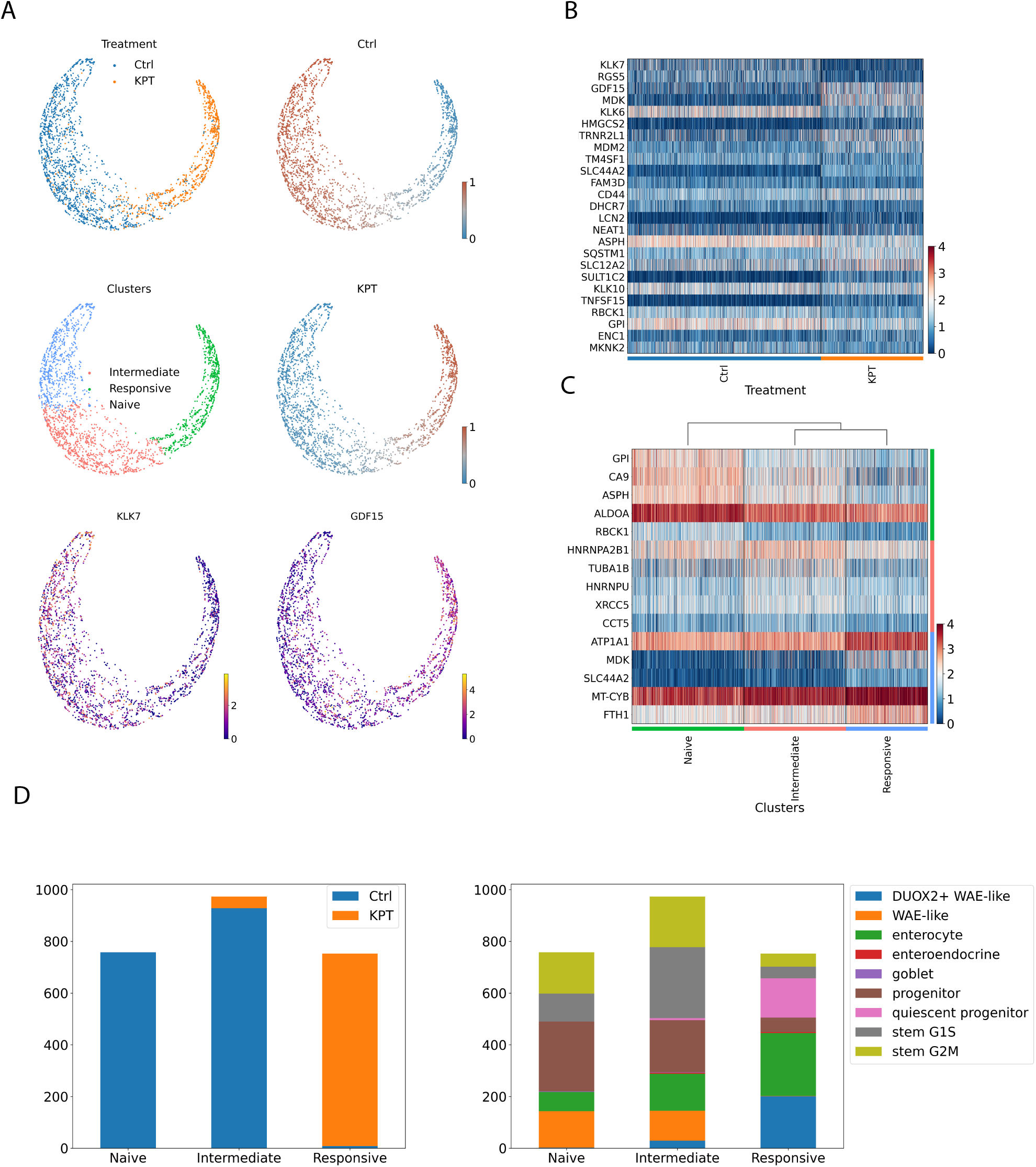
Cellograph identifies treatment groups and distinguishes genes defining these groups on a human organoid dataset. (A) PHATE projection of learned latent space, with cells colored by treatment labels, probabilities of belonging to control or KPT-treated cells, clusters obtained by k-means clustering of the learned latent embedding with *k* = 3, and gene expression of GDF15 and KLK7. (B) Heatmap of top 25 weighted genes from parameterized gene weight matrix. (C) Heatmap of differentially expressed genes between clusters derived from Cellograph. (D) Compositional plot of predicted treatment groups from the softmax probabilities (*p* > 0.5) (left) and cell types annotated by the original study (right) partitioned by clusters.

We also performed k-means clustering on the latent space learned by Cellograph with *k* = 3 (Figure 2A,C). Unlike clustering the original PCA-reduced data, which just focuses on differences in the transcriptome, Cellograph implicitly clusters according to how responsive cells are to the KPT-330 treatment. This successfully groups together cells predicted to belong to the KPT-treated group (called the responsive cluster), a mixed population of cells predicted to be either control or KPT-treated cells (intermediate cluster), and a cluster of cells predicted to be prototypical of the control population (naive cluster).

Based on the softmax probabilities learned by Cellograph, we assigned cells to the control or KPT-treated populations independent of their ground truth labels, and created composition plots according to cluster assignment (fig. 2D). We see that Cellograph’s predictions corroborate the compositional changes in cell types abundance discussed in the original study, namely with decreases in dividing stem cell and progenitor populations, increases in quiescent progenitors, enterocytes, and DUOX2+ WAE-like cells.

Finally, we mapped the cell type annotations onto the clusters obtained by Cellograph (Figure 2D) and observe a high abundance of cycling cells, progenitor cells, and WAE-like cells in the Naive cluster, followed by a decrease of WAE-like cells and progenitor cells in the intermediate population, and a high proportion of DUOX2+ WAE-like cells in the responsive cluster. Altogether, these results demonstrate Cellograph’s ability to identify and visualize cells affected by KPT-330 stimulation.

### 3.2 Cellograph models heterogeneity in cancer drug response during a drug holiday

Encouraged by Cellograph’s performance on the human intestinal organoid dataset, we next investigated how well it could capture heterogeneity in response to cancer drugs under complex treatment regimes. We trained Cellograph on a scRNA-seq dataset of 3042 PC9 cells treated with Erlotinib (Aissa et al., 2021) - a tyrosine kinase inhibitor used to treat non-small cell lung cancer (NSCLC) - for 11 days, followed by withdrawal of the drug for 6 days, referred to as a drug holiday, where select cells were either retreated with Erlotinib or treated with DMSO as a control. The cells were sequenced at 5 timepoints: 0 days with no Erlotinib treatment, 2 days of Erlotinib treatment, 11 days of Erlotinib treatment, at day 19 with or without re-exposure to Erlotinib on day 17, following 6 days of removal from the drug. We trained Cellograph on the dataset with 30 cells labeled for each condition using a 2-layer GCN. We project the learned latent space into 2 dimensions with PHATE, which gives a clear temporal separation of the 6 treatment groups. Coloring cells according to the probability of belonging to each of the conditions provides a narrow distribution of scores in cells in the condition of interest, with the notable exception of Day 11 (Erlotinib before holiday) and Day 19 (Erlotinib after holiday), suggesting a non-uniform response to the drug in these cells both before and after the drug holiday (Figure 3D). The heatmap of the top 25 weighted genes from training (Figure 3B) reveals TUBA1B and CCDC80 as pertinent genes in distinguishing the conditions, which are both markers of drug resistance, with CCDC80 highly expressed in D11 cells, corroborating the original study’s observations of CCDC80, whereas TUBA1B expression in particularly elevated in D19 Erl cells. MT-ND6 is also strongly weighted and appears to uniformly define the population of cells that were treated with DMSO following the drug holiday. This is a mitochondrial gene which has been previously implicated in colorectal adenocarcinoma and associated with changing energy requirements due to cells aggressively proliferating (Wallace et al., 2016). Clustering the learned latent space identifies three clusters among these two conditions (Figure 3A,D), one consisting of cells predicted to have a prototypical response after 11 days of Erlotinib treatment (cluster 3), and similarly for day 19 after re-exposure to the drug (cluster 5), followed by a mixed population of both cell types (cluster 2). Differential expression between the three clusters (Figure 3C) identified high expression of TUBA1B in cluster 5, which is associated with poor prognosis in NSCLC, suggesting persisting drug tolerance after the holiday period. Similarly, we observe differential expression of INHBA in cluster 3, a senescence mediator that’s associated with prognosis in many cancer types (Zhao et al., 2022). Interestingly, TUBA1B and INHBA expression are significantly reduced in the day 19 population that was not retreated with Erlotinib. Altogether, Cellograph captures clinically relevant genes driving heterogeneity in response to treatment, as well as identifying additional genes such as MT-ND6 that were not captured using previous approaches.

**Figure 3:**
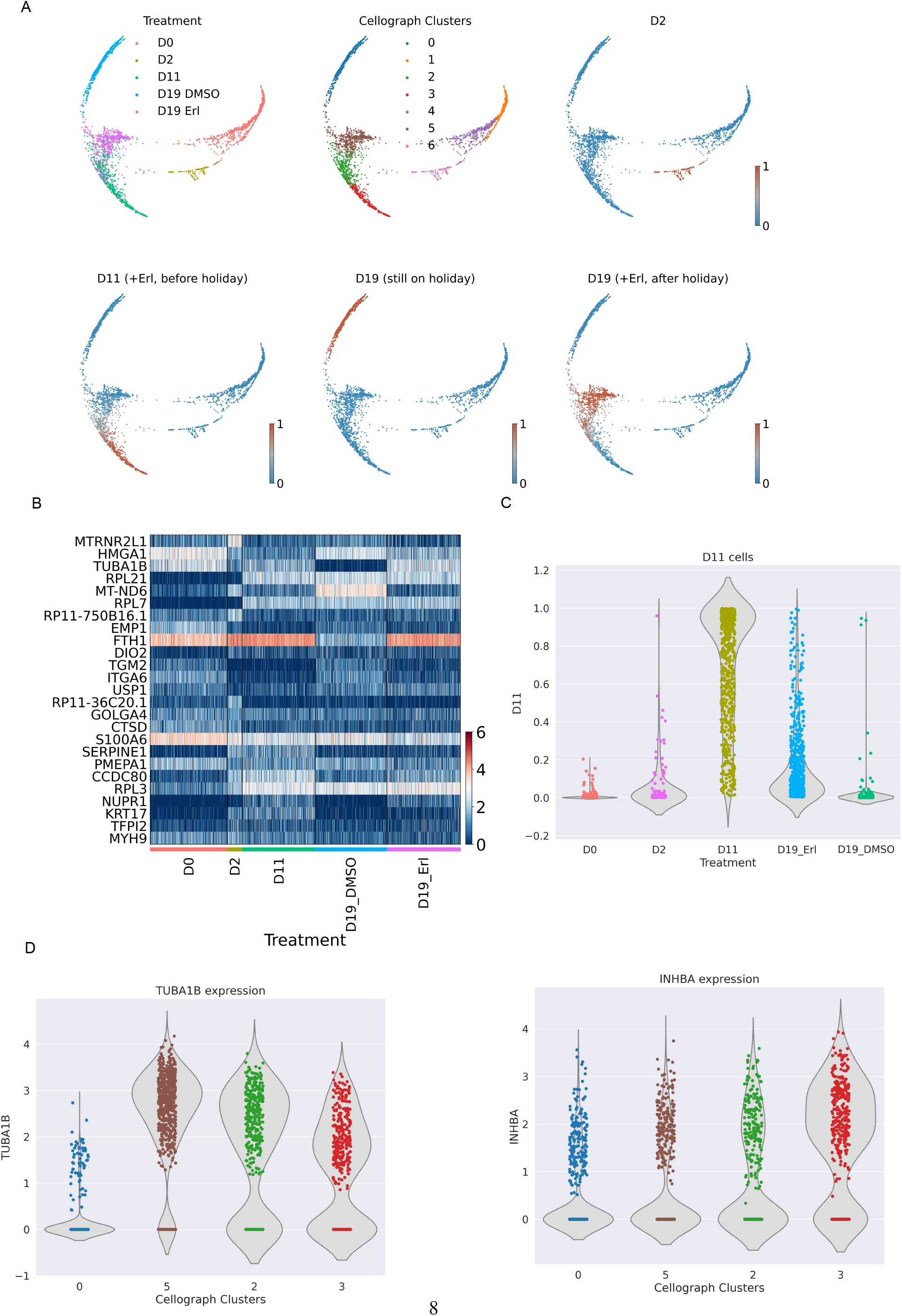
Cellograph defines genetic signatures of distinct drug responses in the drug holiday dataset. (A) PHATE embeddings of the learned latent space colored according to the treatment labels, clusters, and treatment probabilities (day 0 not shown). (B) Heatmap of top 25 weighted genes from learned parameterized gene weight matrix. (C) The distribution of treatment probabilities for Day 11 cells partitioned by treatment groups. (D) The distribution of gene expression between clusters 0, 5, 3, and 2 of select differentially expressed genes (INHBA, TUBA1B).

### 3.3 Cellograph distinguishes between transdifferentiation and dedifferentiation in myogenesis

Finally, we assessed Cellograph’s ability to distinguish cells undergoing distinct cell state transitions temporally on a scRNA-seq dataset of 33,380 mouse embryonic fibroblasts (MEF) undergoing either dedifferentiation to adult stem cells called induced myogeneic progenitor cells (iMPCs) or myogenic transdifferentiation to myotubes (Yagi et al., 2021). The original study was motivated to understand the transcriptional and epigenetic mechanisms of how over-expression of the MyoD transcription factor induced MEFs to undergo reprogramming to either myotubes or iMPSCs with a MyoD-inducible transgenic model. The myotubes were induced by overexpression of MyoD, while the addition of small molecules produced Pax7^+^iMPSCs that were very similar to primary muscle stem cells. The authors used trajectory analysis via diffusion maps and UMAP embeddings of combined single-cell data of MEFs expressing MyoD or MyoD + a cocktail of small molecule inhibitors (forskolin, RepSox, and CHIR99021, collectively abbreviated as “FRC” in the original paper) to reveal that dedifferentiation and transdifferentiation follow two different trajectories.

We trained Cellograph on this dataset with 200 cells per treatment group labeled for training for 400 epochs and obtained a single trajectory that starts with transdifferentiation and culminates in dedeifferentiation to Pax7^+^iMPCS. The original study revealed an overlap between the major fraction of day 4 MyoD-treated cells and day4/8 MyoD+FRC-treated cells in their UMAP and DPT embeddings. Interestingly, however, Cellograph detects no significant overlap (Figure 4A), which is further supported by the derived probabilities of belonging to each of the experimental groups. Looking at the top-weighted genes from training the model, high expression of CRABP1 and LUM distinguished the transdifferentiating population, whereas dedifferentiation was weighted by high expression of cyclin D1, suggesting cell cycle entry is a necessary step to producing iMPCs. CRABP1 is known to promote stem cell proliferation by its downregulation (Lin et al., 2017). However, it does not appear to inversely vary with cyclin D1 nor directly vary with CDKN1C, another top-weighted gene associated with dedifferentiation that codes for p57 (Matsuoka et al., 1995), a well-characterized cell cycle inhibitor, thus making it an interesting candidate gene to investigate further for its functional role in myogenesis. Furthermore, CDKN1C is known to be stimulated by MyoD expression (Figliola and Maione, 2004). Interestingly, Pax7^+^iMPCs have minimal cyclin D1 expression and instead high expression of CDKN1C. Satellite cells re-enter the cell cycle in response to injury through asymmetric division, thus, this cell population likely resides in a quiescent, unstimulated state.

**Figure 4:**
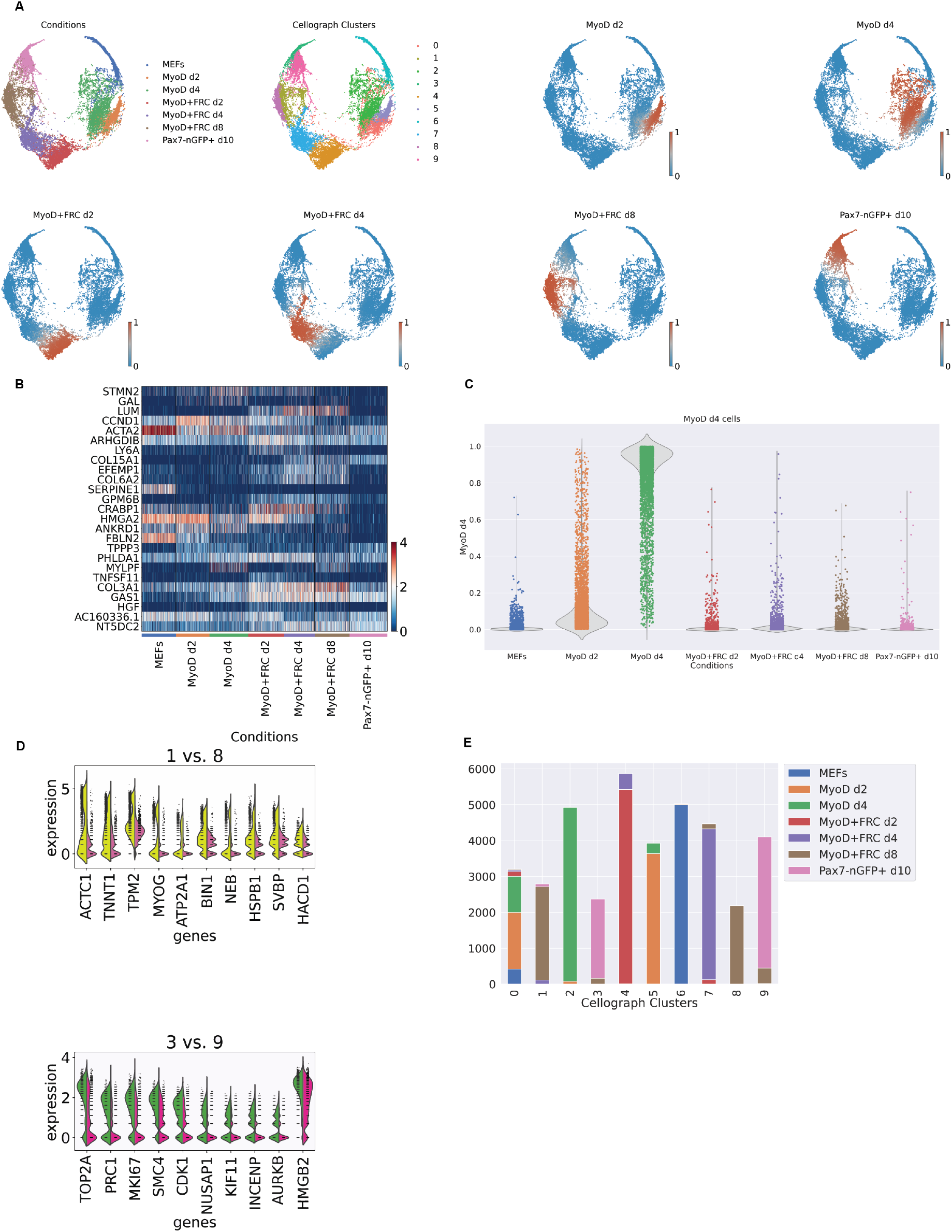
Cellograph distinguishes the molecular mechanisms of transdifferentiation and dedifferentiation in myogenesis. (A) PHATE embeddings of learned latent space annotated according to treatment conditions, clusters, and softmax probabilities of all conditions except for MEFs, defining the in-group variation. (B) Heatmap of top weighted genes from parameterized gene weight matrix, identifying pertinent genes such as cyclin D1 and CRABP1. (C) Violin plot of softmax probabilities of cells belonging to the MyoD/day 4 treatment group, showing similarities to the MyoD/day 2 population. (D) Violin plots of top 20 differentially expressed genes between clusters 1 and 8 and clusters 3 and 9, which define the Pax7^+^cells and MyoD+FRC/day 8 treated cells, respectively. (E) Compositional plot of predicted cell types partitioned by cluster

STMN2, an early neuronal marker, was also identified as a pertinent gene in distinguishing between these processes, with high expression in the transdifferentiation condition, perhaps owing to the instability and inefficiency of generating myotubes with MyoD alone. Other genes exhibit more monotonically changing trends, such as GM4218, PRUNE2, and TIMP1, that correspond to the silencing of the MEF program. Clustering the latent space and mapping the clusters onto the PHATE embedding distinguished between the different treatment conditions and heterogeneity in the MyoD+FRC day 8 condition, in particular, differential expression of MYOG, which specifies the myotube fate, was observed in the majority of cells, and corroborates observations from trajectory analysis in the original study, where this gene is observed in both trajectories. Altogether, Cellograph is able to successfully distinguish these biological processes, and identify additional gene programs explaining these differences.

### 3.4 Cellograph outperforms published differential abundance methods and popular single-cell clustering methods

Finally, we benchmarked Cellograph’s performance in identifying cells most impacted by perturbations against three published methods: MELD, Milo (Dann et al. (2021)), and Covarying Neighbor Analysis (CNA) (Reshef et al. (2021)) - and compared its clustering performance to popular single-cell clustering methods. We used the categorical crossentropy objective function in the first evaluation as we believed a method quantifying experimental perturbations should capture a broad range of signals for each experimental label it is trying to predict. In particular, it quantifies the difference between two pairs of discrete probability distributions by calculating the following sum,

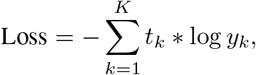

where *t_k_* corresponds to a ground truth label and *y_k_* is the softmax probability for the kth class. It is commonly used in classification tasks for neural networks as it favors high probabilities for stereotypically correct assignments while low probabilities for others, which motivated us to use it as our objective function in training Cellograph. When applied to all the cells in our datasets, we obtain consistently lower loss scores than MELD (Table 1).

**Table 1:** Categorical cross-entropy of Cellograph vs MELD on all cells

| Dataset | Cellograph | MELD (optimal settings) |
| --- | --- | --- |
| Organoid | 0.297 | 0.374 |
| Drug Holiday | 0.252 | 0.452 |
| Myogenesis | 0.193 | 0.605 |

When evaluating Cellograph relative to Milo and CNA, however, we could not perform direct quantitative comparison. Milo tests for differential abundance on kNN graphs by aggregating cells into overlapping neighborhoods and performing a quasi-likelihood F test. This returns a metric of the log-fold change of the differential abundance in each neighbor, not a single-cell measurement giving the probability of that cell belonging to one treatment class versus another. Thus, we cannot perform a direct quantitative comparison and instead present a qualitative assessment of performance. Running Milo on the human organoid dataset, we observe a positive correlation between the Milo-derived log-fold changes in differential abundance and the probability of cells belonging to the KPT-treated group. However, when applied to the the drug holiday and myogenesis datasets, which have more complex experimental designs with multiple conditions, Milo fails to yield clear, interpretable results, with low DA in the untreated population, high DA in the cells after one day of Erlotinib treatment, and minimal DA in all other conditions. Similarly, in the myogenesis dataset, we observe high DA in Pax7-treated cells, low DA in MEFs, and minimal DA everywhere else.

CNA performs association analysis agnostic of parameter tuning, making it an efficient method. Like Milo, it aggregates cells into neighborhoods, and calculates a neighborhood abundance matrix (NAM), where each entry *C_n_*, *m* is the relative abundance of cells from sample n in neighborhood m. From there, it derives principal components where positive loadings correspond to higher abundance while negative loadings correspond to lower abundance. This enables the characterization of transcriptional changes corresponding to maximal variation in neighborhood abundance across samples. Association testing is performed between transcriptional changes and attributes of interest using the first *k* NAM-PCs. It returns the Spearman correlation between the attribute and abundance of the neighborhood anchored at each cell. Applied to the human organoid dataset with the KPT treatment status as the attribute of interest, we obtain similar results as our method, MELD, and Milo with high correlation observed in the KPT-treated cells, and low correlation in the observed cells, compared to the probability of observing cells in the KPT-treated group. However, on the Erlotinib and myogenesis datasets, like Milo, we obtain results incongruous with Cellograph or MELD’s performance, or even Milo. High abundance is predicted for cells treated right before holiday and following the holiday, regardless of whether cells were retreated with Erlotinib, whereas low abundance is observed in both the untreated cells and cells treated with Erlotinib for one day, while cells with 11 days of treatment have zero correlation. Since this dataset spans multiple conditions whereas unlike MELD or Cellograph, CNA calculates one set of metrics, it was difficult to interpret these results in the context of the experiment. We obtained similarly incongruous results for the myogenesis dataset (Figure 5). Altogether, Cellograph provides robust and interpretable results for more complex experimental designs with multiple treatment groups compared to CNA and Milo, and performs consistently better than MELD with a significantly lower runtime for optimal performance (Figure 6).

**Figure 5:**
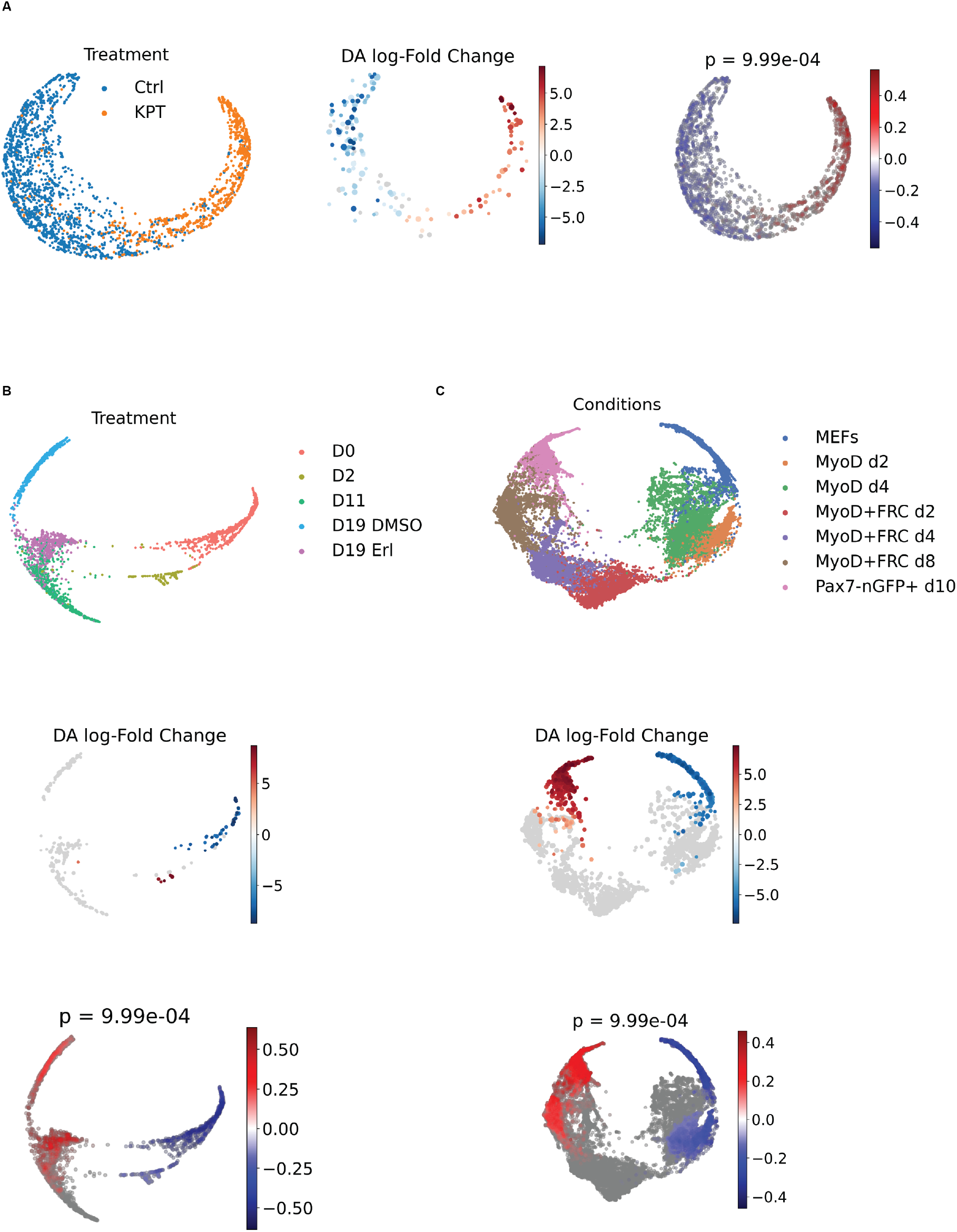
Results of running Milo and CNA on the datasets evaluated. (A) Output of running Milo and CNA on the human organoid dataset. (B) Output of running Milo and CNA on the drug holiday dataset. (C) Output of running Milo and CNA on the myogenesis dataset

**Figure 6:**
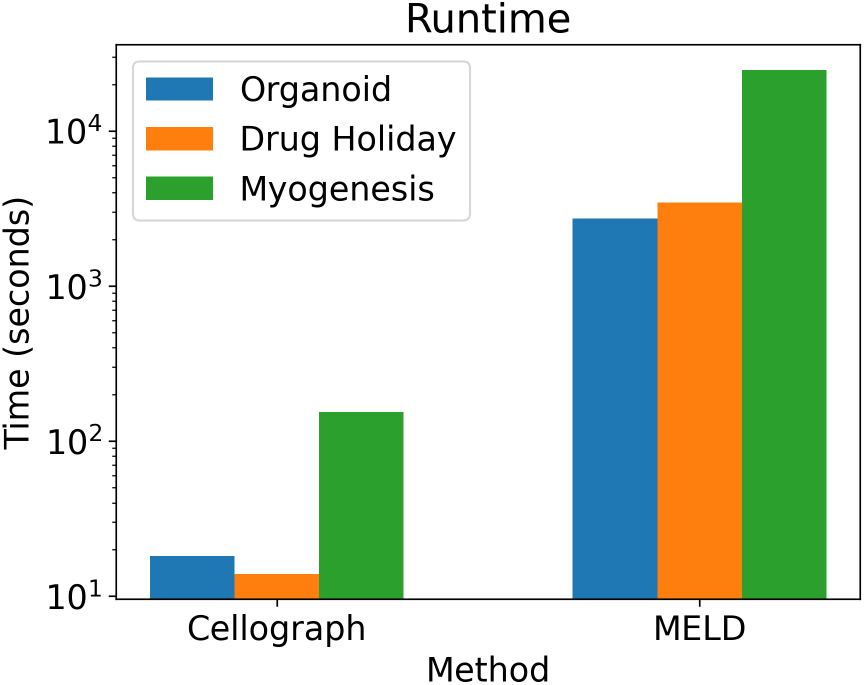
Runtime of Cellograph’s performance versus MELD’s on optimal parameter settings under each of the datasets on a log scale. Cellograph consistently outperforms MELD on each dataset, while using fewer computing resources.

When evaluating clustering performance with NMI, *k*-means clustering on the learned latent space consistently outperformed popular algorithms, Leiden, Louvain, and *k*-means clustering on data in PCA space (100 independent runs for each clustering approach) (Figure 7). Altogether, Cellograph outperforms MELD is estimating how prototypical cells are of their ground truth labels, and consistently ranks higher than standard algorithms for clustering.

**Figure 7:**
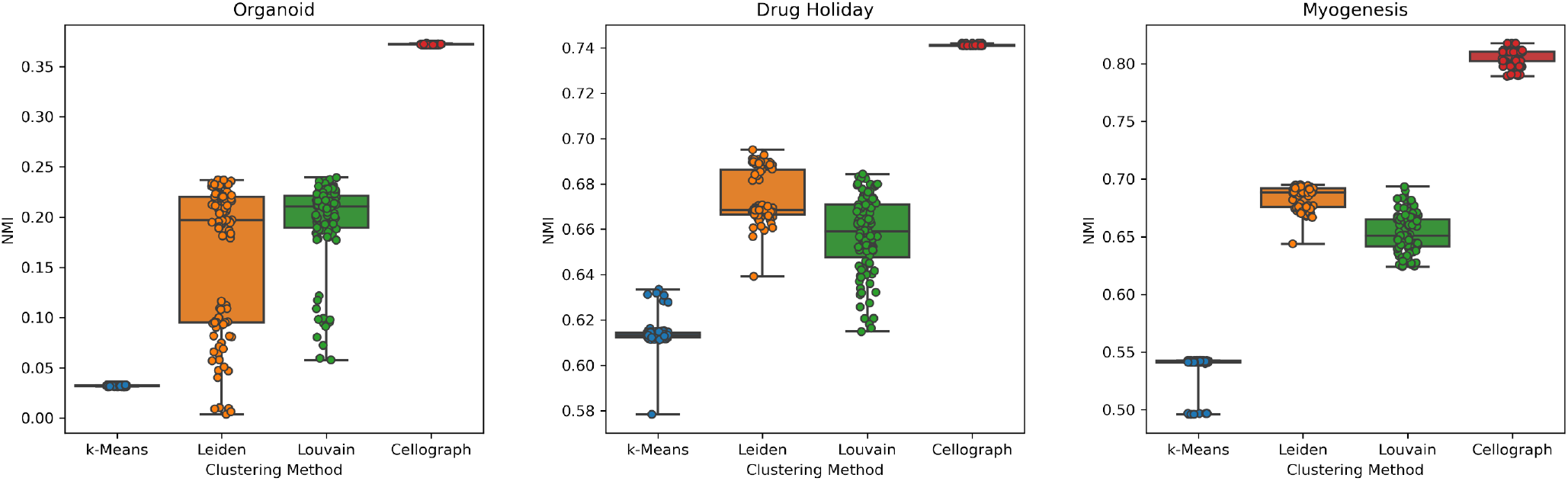
Distributions of 100 independent NMI calculations for each clustering algorithm for all three datasets evaluated, quantifying concordance between the cluster assignments and ground truth labels

## 4 Discussion

When designing single-cell experiments exploring the impacts of different treatments, it is vital to leverage the single-cell heterogeneity present. The increasing complexity of the experimental design (e.g., multiple treatments, various timepoints, etc) can result in diminishing returns from standard differential gene expression and clustering approaches due to the biological and technical variability present at the single-cell level. Existing approaches like MELD and VFC are apt for studying the effects of one experimental treatment, but cannot be easily generalized to more complicated experimental programs. We designed Cellograph to address this challenge. Beyond just quantifying single-cell responses to perturbations analogously to MELD, Cellograph’s primary innovation lies in its novel way of visualizing and clustering single-cell data by means of graph neural networks, which, through the parameterized gene weight matrix, provides its interpretable means of understanding which genes drive the difference between conditions. We have shown that our approach improves clustering on three diverse datasets compared to standard clustering approaches, as well as captures a stronger signal of the ground truth experimental label compared to MELD. Clustering agnostic of experimental conditions can fail to take into consideration the diversity of cellular responses to these perturbations and how those correspond to the transcriptomic variation. By applying simple k-means clustering to the latent space, we can obtain more informative clusters that enable deeper biological insight, especially in populations under the same experimental treatment. In addition to improved differential gene expression, we also obtain complementary information from the parameterized weight matrix after training, which reveals the most important genes in distinguishing between different treatments.

In a published dataset of donor-provided organoid samples, we were able to successfully corroborate original findings, while providing a visually informative view of the data, and revealed novel insights into drivers of KPT-mediated organoid differentation. Similarly, in our drug holiday application, we identified additional markers of drug resistance using the parameterized gene weight matrix, and described heterogeneity of cells in response to Erlotinib after 11 days and post-holiday, while characterizing the popular that was retreated after the holiday that could inform future experiments into druggable targts for NSCLC. Finally, in our myogenesis evaluation, we identified shared features between transdifferentiation and dedifferentiation, while capturing relevant markers that distinguished the two processes. We anticipate Cellograph will find a wide range of application to other biological contexts and different single-cell modalities as an all-in-one framework for facilitating visualization, clustering, and single-cell responses to perturbations, on top of its efficiency. For example, this work could find utility in clinical applications to studying heterogeneity in patient-treated samples in response to an experimental cancer drug. This could be also employed to study impacts of cancer drugs on cell cycle in protein immunofluorescence imaging data (Gut et al., 2018). The graph neural network architecture of node classification could even be extended to graph classification for looking at multiple patient samples, as is common in mass cytometry, or regression to predict continuous variables such as cellular pseudotime in the context of differentiation, cell cycle age (Stallaert et al., 2022), or gestational age in data from pregnant women (Aghaeepour et al., 2017), and may be further explored in future work.

